# Ectonucleotidases and Purinergic Receptors in Mouse Prostate Gland

**DOI:** 10.1101/2024.09.30.615810

**Authors:** Jovian Yu, Christina Sharkey, Aria Olumi, Zongwei Wang

## Abstract

Extracellular ATP/ADP and its metabolite adenosine are important signaling molecules that regulate cellular function by binding to P2 and P1/adenosine receptors. The kinetics of these signaling molecules are critically modulated by ectonucleotidases, enzymes that convert ATP/ADP to adenosine. Although the expression and function of these enzymes and relevant purinergic receptors in the prostate gland are not well understood, recent reports indicate impaired ATP hydrolysis activity in the aging prostate. Purinergic signaling is known for its role in inflammation, muscle contraction, pain sensation, and cell proliferation in many systems, suggesting its potential importance in normal prostate function and pathological conditions such as benign prostatic hyperplasia and prostatitis. To better understand purine-converting enzymes and purinergic receptors in the prostate, we isolated mouse prostate glands for immunofluorescent staining and microscopy imaging using specific antibodies. Our study identified a differential expression profile of purinergic enzymes and receptors in the prostate: ENTPD1 and P2×1 receptors predominantly in prostate smooth muscle cells, ENTPD2 and NT5E in prostate interstitial cells, and ALPL in prostate epithelial cells. Functionally, in addition to the P2×1-mediated prostate smooth muscle contraction induced by agonist α,β-meATP, we observed an ATPγS-induced contraction force after P2×1 desensitization. This led to the identification of multiple P2Y receptors in mouse prostate smooth muscle, including P2Y1, P2Y2, and P2Y11 receptors, which potentially mediate the ATPγS-induced contraction force. These discoveries lay the foundation for further mechanistic understanding of how purinergic signaling regulates prostate function and dysfunction in both rodents and potentially humans.

## Introduction

The prostate gland, a male organ composed of tubular structures, plays a crucial role in fertility. Luminal epithelial cells secrete fluids that mix with sperm to create semen, while stromal smooth muscle cells contract forcefully to expel semen into the urethra during ejaculation[1]. Dysregulation of prostate function can lead to benign prostatic hyperplasia (BPH), prostatitis, and even cancer, which are significant morbidities in the aging population[2]. For instance, BPH affects 50% of men over 50 and >80% of men over 70. However, the molecular physiology of the prostate gland and the pathogenesis of its disorders are still not fully understood, necessitating a deeper understanding of the underlying mechanisms for improved therapeutic strategies[3].

Purinergic signaling plays numerous important roles, including in inflammation and immune responses, muscle contractility, gluconeogenesis and metabolism, cell proliferation and differentiation, tumor metastasis, and cancer therapy[4]. This signaling comprises a dynamic and complex interactive network. Cell-released signaling molecules like ATP and ADP bind to P2 purinergic receptors of the P2X (1-7) and P2Y (1,2,4,6,11-14) families, initiating downstream signaling. ATP and ADP are further converted into adenosine by enzymes. Ectonucleoside triphosphate diphosphohydrolases (ENTPD1, 2, 3, 8), a group of cell surface ectonucleotidases sequentially convert extracellular ATP to ADP and then to AMP, while ecto-5’-nucleotidase (NT5E) and alkaline phosphatase (ALPL) convert AMP to adenosine. Adenosine binds to P1 or adenosine receptors (A1, A2a, A2b, A3)[5].

Physiologically, purinergic contractility accounts for part of nerve-mediated prostate contraction, although the detailed mechanism of how this signaling network impacts prostate gland function is not fully understood[6; 7; 8; 9; 10]. However, abnormal ATP levels and purinergic receptor function have been reported in both animal models and human patients with prostate dysfunctions. For example, in rodent models of prostatitis, elevated levels of P2×2, P2×3, and P2×7 might mediate prostatic pain sensation[11; 12]. Increased expression of P2×1 in aged mice could cause an increase in prostate muscle tone[9; 13]. In human BPH patients, increased ATP release and impaired ATP hydrolysis have been observed, along with a significant reduction in P2×2 and P2×3 proteins[13]. These studies suggest that purinergic signaling in the prostate gland plays an important role in prostate physiology and pathophysiology.

Our laboratory investigates molecular pathways leading to BPH, focusing on the regulation of steroid 5-alpha reductase in androgen and prostate growth[14; 15; 16; 17; 18]. In this study, we broaden our research by determining various purine-converting enzymes and purinergic receptors in the mouse prostate gland. Additionally, we have identified a novel purinergic contractility, potentially mediated by P2Y receptor(s) in the smooth muscle of the mouse prostate. These findings provide a foundation for further mechanistic insights into how purinergic signaling regulates prostate function and dysfunction.

## Materials and Method

### Materials

Unless otherwise specified, all chemicals were obtained from Sigma (St. Louis, MO) and were of reagent grade or higher. α,β-meATP (Cat. #: 3209) and NF 546 (Cat. #: 3892) were purchased from R&D system (Minneapolis, MN, USA). ATPγS (Cat. #: NU-406) and ADPβS (Cat. #: NU-433) were obtained from Jena Bioscience (Thuringia, Germany).

### Animals

Male C57BL/6J mice (Jackson Laboratory, Bar Harbor, ME) aged 12-16 weeks old were used in this study. All animal studies were performed in adherence with U.S. National Institutes of Health guidelines for animal care and use and with the approval of the Beth Israel Deaconess Medical Center Institutional Animal Care and Use Committee (Protocol #: 2022-049). Mice were housed in standard polycarbonate cages and maintained on a 12:12-h light-dark cycle at 25°C with free access to food and water.

### Isolation of mouse prostate glands

A midline abdomen incision was made to expose and remove the entire prostate along with the seminal vesicle, bladder, urethra, and vas deferens. The tissue was placed in a petri dish with PBS solution, and the prostate was carefully dissected out under an Olympus dissecting microscope. The isolated prostate tissue was then used for either myography study or immunofluorescence staining.

### Immunofluorescence staining and microscopy imaging

The isolated whole prostate gland was fixed in 4% (wt/vol) paraformaldehyde for 2 hours at room temperature. The fixed tissue was then treated with 30% (w/v) sucrose solution for another 2 hours, cryoprotected, and frozen in OCT compound at -80 °C. The tissue was sectioned (5 μm) and incubated with primary antibodies (1:100) overnight at 4°C. Depending on the number and species of the primary antibodies, the sections were then incubated with Alexa Fluor 488– and/or Alexa Fluor 555–conjugated secondary antibodies (diluted 1:100), and nuclei were stained with DAPI. Imaging was performed on an Olympus BX60 fluorescence microscope with a 40×/0.75 objective. Images (512 and 512 pixels) were saved as TIFF files and imported into Adobe Illustrator 28.1. In this study, each prostate tissue was sectioned to obtain slides with approximately four sections of tissue per slide. Each antibody staining was repeated at least twice to ensure consistency of the staining results.

### Antibodies

Detailed information on antibodies and dyes used for immunofluorescent staining is listed in Table 1.

**Table 1:** List of antibodies and dyes used for immunofluorescent staining.

| Antibody or dye | Catalog # | Host animal | Company |
| --- | --- | --- | --- |
| ENTPD1 | AF4398 | Sheep | R&D System |
| P2X1 | APR-001 | Rabbit | Alomone Lab |
| ENTPD2 | AF5797 | Sheep | R&D System |
| CD34 | AB8158 | Rat | ABCAM |
| ENTPD3 | AF4464 | Sheep | R&D System |
| NT5E | AF4488 | Sheep | R&D System |
| ALPL | AF2910 | Goat | R&D System |
| P2Y1 | H120 | Rabbit | Santa Cruz |
| P2Y2 | AB10270 | Rabbit | ABCAM |
| P2Y11 | A24482 | Rabbit | ABclonal |
| Alexa 488 Donkey anti-Sheep | A-11015 | Donkey | ThermoFisher |
| Alexa 488 Donkey anti-Rabbit | A-21206 | Donkey | ThermoFisher |
| Alexa 555 Donkey anti-Rat | A-78945 | Donkey | ThermoFisher |
| Alexa 488 Donkey anti-Goat | A-11055 | Donkey | ThermoFisher |
| Rhodamine Phalloidin | PHDR1 |  | Cytoskeleton |
| Antifade mounting medium with DAPI | H-2000-10 |  | Vector Laboratories |

### Myography

Each male mouse contains two long anterior prostate lobes, and a longitudinal middle cut of the prostate yields two tissue preparations for myograph studies, with one suture tied to the tip of the anterior prostate lobe and the other suture tied to the dorsal lobe end. Prostate strips were then mounted in an SI-MB4 tissue bath system (World Precision Instruments, Sarasota, FL, USA). Force sensors were connected to a TBM 4M transbridge (World Precision Instruments), and the signal was amplified by a PowerLab (AD Instruments, Colorado Springs, CO, USA) and monitored through Chart software (AD Instruments). Prostate strips were gently stretched to optimize contraction force, then pre-equilibrated for at least 1 h. All experiments were conducted at 37°C in physiological saline solution (in mM: 120 NaCl, 5.9 KCl, 1.2 MgCl_2_, 15.5 NaHCO_3_, 1.2 NaH_2_PO_4_, 11.5 glucose, and 2.5 mM CaCl_2_) with continuous bubbling of 95% O_2_ / 5% CO_2_. Contraction force was sampled at 2000/s using Chart software. Prostate strips were treated with agonists or antagonists, and/or subjected to electrical field stimulation (EFS).

### Electrical field stimulation

Prostate strip EFS was carried out by a Grass S48 field stimulator (Grass Technologies, RI, USA) using standard protocols previously described: voltage 50 V, duration 0.05 ms, trains of stimuli 3 s, and frequencies 20, and 50 Hz [19; 20].

### Statistical analyses

All data are expressed as means ± SD, or presented as boxes and whiskers (extending from minimum to maximum values). Data were analyzed by Student’s *t*-test between two groups. P < 0.05 was considered significant.

## Results

### ENTPD1 and P2×1 receptors are predominantly expressed in prostate smooth muscle cells

ENTPD1 is a major enzyme that converts ATP and ADP to AMP. Immunostaining and imaging have demonstrated the expression of ENTPD1 in the prostate gland (Figure 1A). By counterstaining the actin filament with rhodamine-phalloidin and nuclei with DAPI, we can identify its cellular localization clearly. As shown in Figure 1B, rhodamine-phalloidin binds to actin-rich smooth muscle cells, indicated by a strong bright red signal. In contrast, epithelial cells, aligned inside the muscle bundles, exhibit relatively large, round nuclei with faint red actin signaling outlining their membranes. ENTPD1 expression is predominantly found in the smooth muscle bundles of the prostate gland. It is also strongly expressed in a subset of cells in the interstitial space, likely the endothelial cells of the vasculature, as previously reported in other systems[21; 22]. The P2×1 receptor, an ATP-activated cation channel, is solely expressed in smooth muscle cells in the prostate gland, consistent with previous reports that the P2×1 receptor mediates prostate contractility[7; 8; 9].

**Figure 1.**
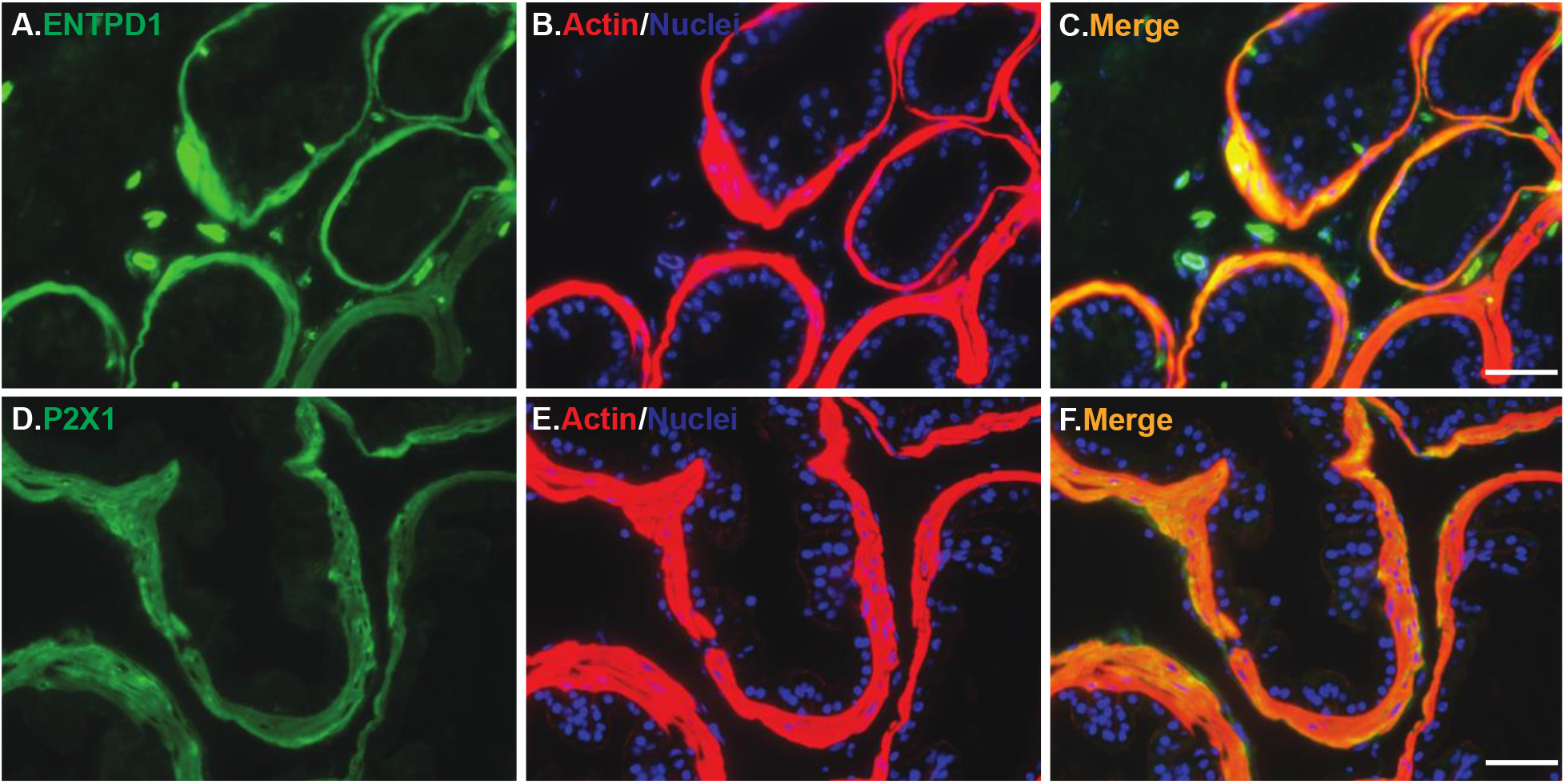
Expression of ENTPD1 and P2×1 receptors in the smooth muscle of mouse prostate. Mouse prostate tissue was labeled with specific anti-ENTPD1 (green in panel A) and anti-P2×1 (green in panel D) antibodies. These antibodies colocalized with prostate smooth muscle (red in panels B and E) as shown in the merged images (yellow in panels C and F). Nuclei were stained with DAPI (blue). The white scale bar in the bottom right corner of panels C and F represents 50 μm.

### ENTPD2 but not ENTPD3 is expressed in prostate interstitial cells

ENTPD2 is another major purine-converting enzyme located on the extracellular surface. As shown in Figure 2A-C, ENTPD2 expression in the prostate gland is not colocalized with smooth muscle cells or epithelial cells. Instead, it is found in cells with long, thin filaments adjacent to the smooth muscles. To identity these cells, we labeled them with the interstitial cell marker CD34. Figure 2D-F demonstrate that ENTPD2 is nicely colocalized with CD34 signaling, indicating that these ENTPD2-positive cells are interstitial cells in the prostate gland.

**Figure 2.**
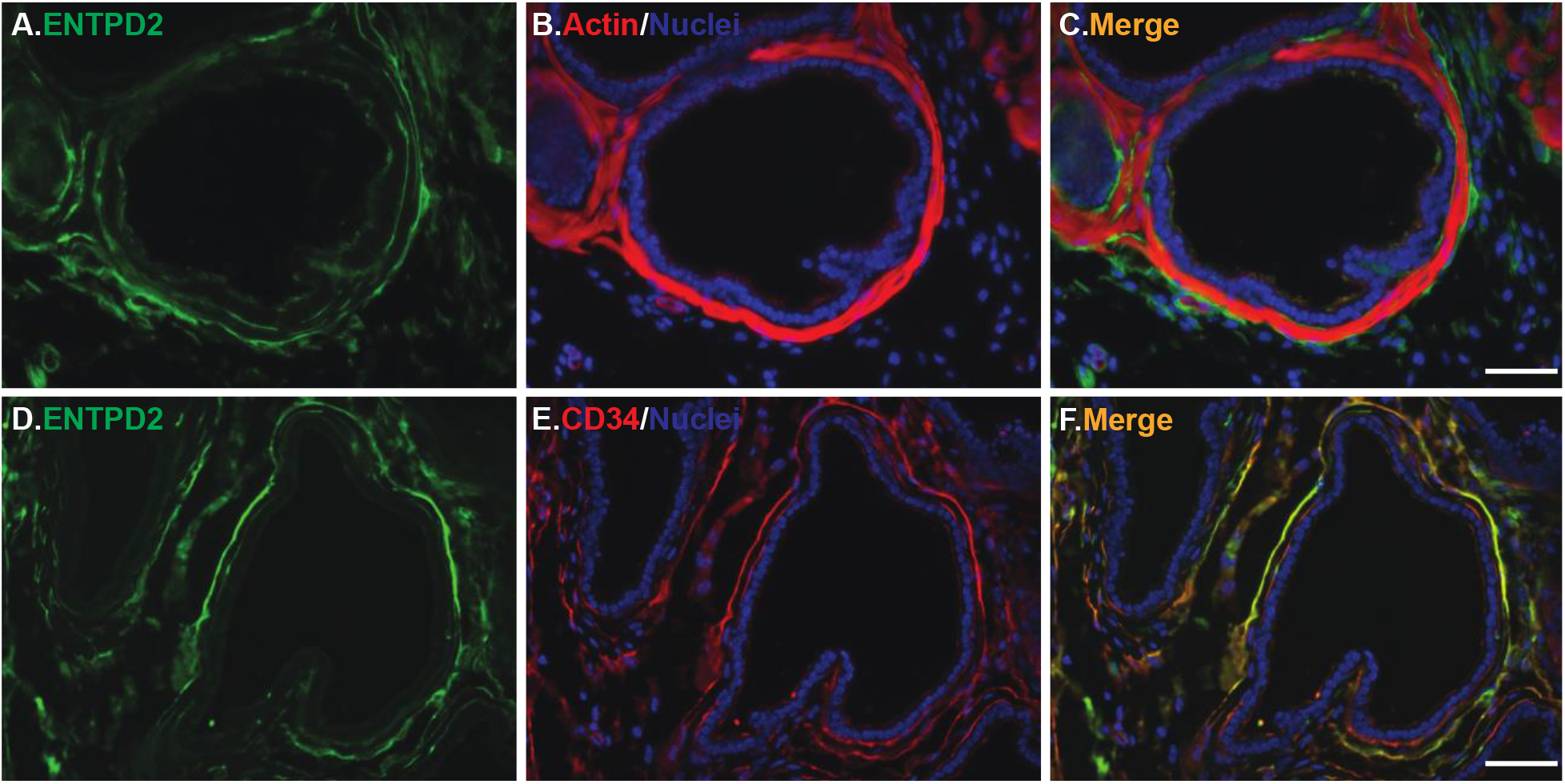
Expression of ENTPD2 in mouse prostate interstitial cells. Mouse prostate tissue was labeled with specific anti-ENTPD2 (green in panels A and D) antibody. ENTPD2 did not colocalize with prostate smooth muscle (red in panels B), as shown in the merged images (panel C). However, it colocalized with the interstitial cell marker CD34 (red in panel E), as shown in the merged images (yellow in panel F). Nuclei were stained with DAPI (blue). The white scale bar in the bottom right corner of panels C and F represents 50 μm.

ENTPD3, another enzyme with a similar function, is expressed in the basolateral membrane of bladder epithelial cells (Figure 3D-F). However, ENTPD3 expression was not detected in any cell type within the prostate gland (Figure 3A-C).

**Figure 3.**
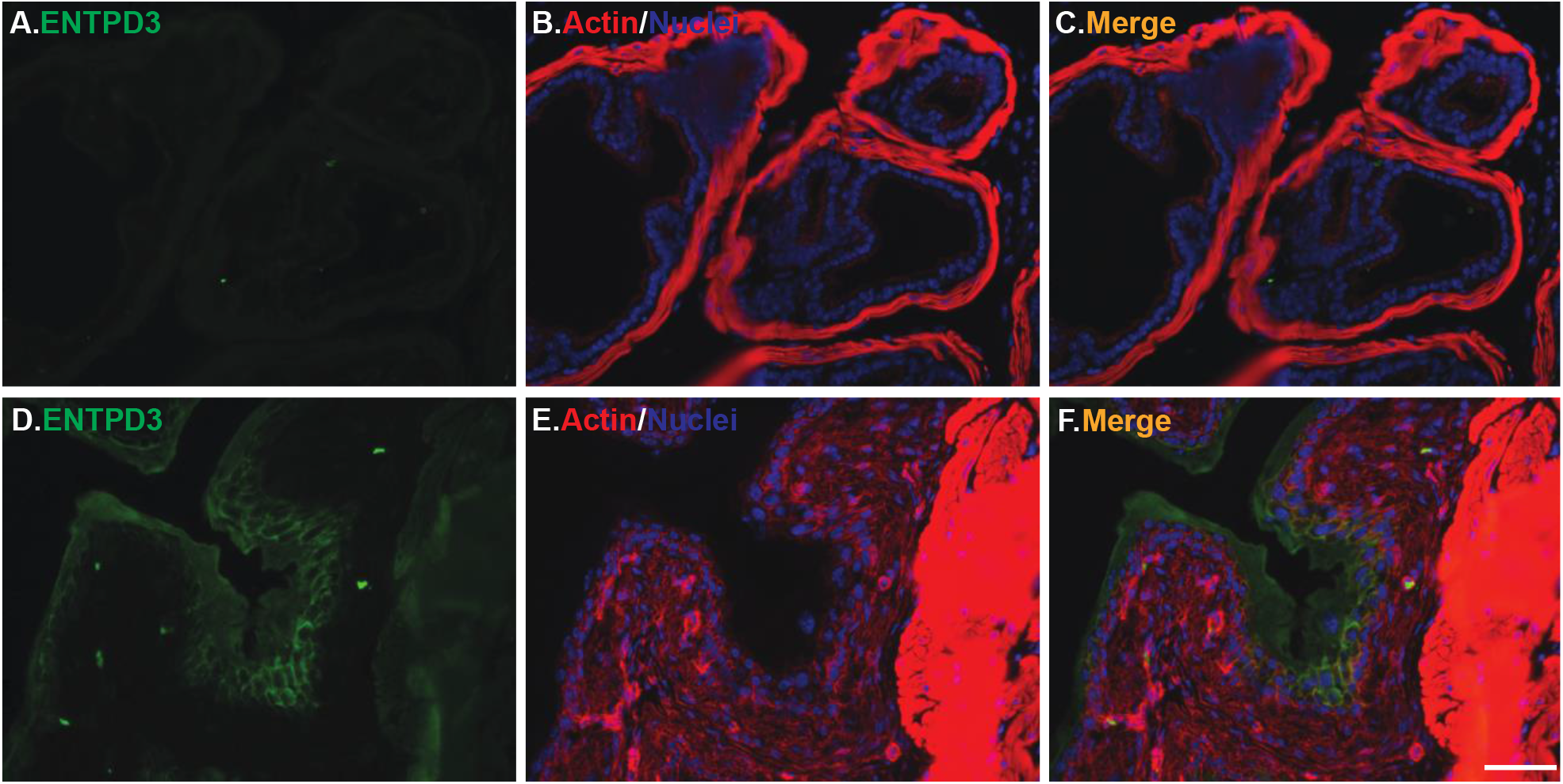
Absence of ENTPD3 in mouse prostate gland. Mouse prostate and bladder tissues were labeled with specific anti-ENTPD3 antibody (green in panels A and D). ENTPD3 was localized on the basolateral membrane of bladder urothelial cells (green in panel D, serving as a positive control), but it was not observed in prostate epithelial cells. Merged images are shown in panels C and F. Nuclei were stained with DAPI (blue). The white scale bar in the bottom right corner of panels C and F represents 50 μm.

### NT5E and ALPL are differentially expressed in the prostate gland

NT5E is an enzyme that specifically converts AMP to adenosine, while ALPL is a non-specific enzyme converting ATP/ADP to AMP and adenosine. Interestingly, NT5E is only present in cells morphologically similar to ENTPD2-positive cells, but not in smooth muscle or epithelial cells (Figure 4A-C), indicating its expression in prostate interstitial cells. In contrast, ALPL is strongly expressed in the prostate epithelial cell membrane, particularly on the apical membrane surface (Figure 4D-F), suggesting an important role for ALPL in converting luminal ATP/ADP molecules into adenosine in the prostate gland.

**Figure 4.**
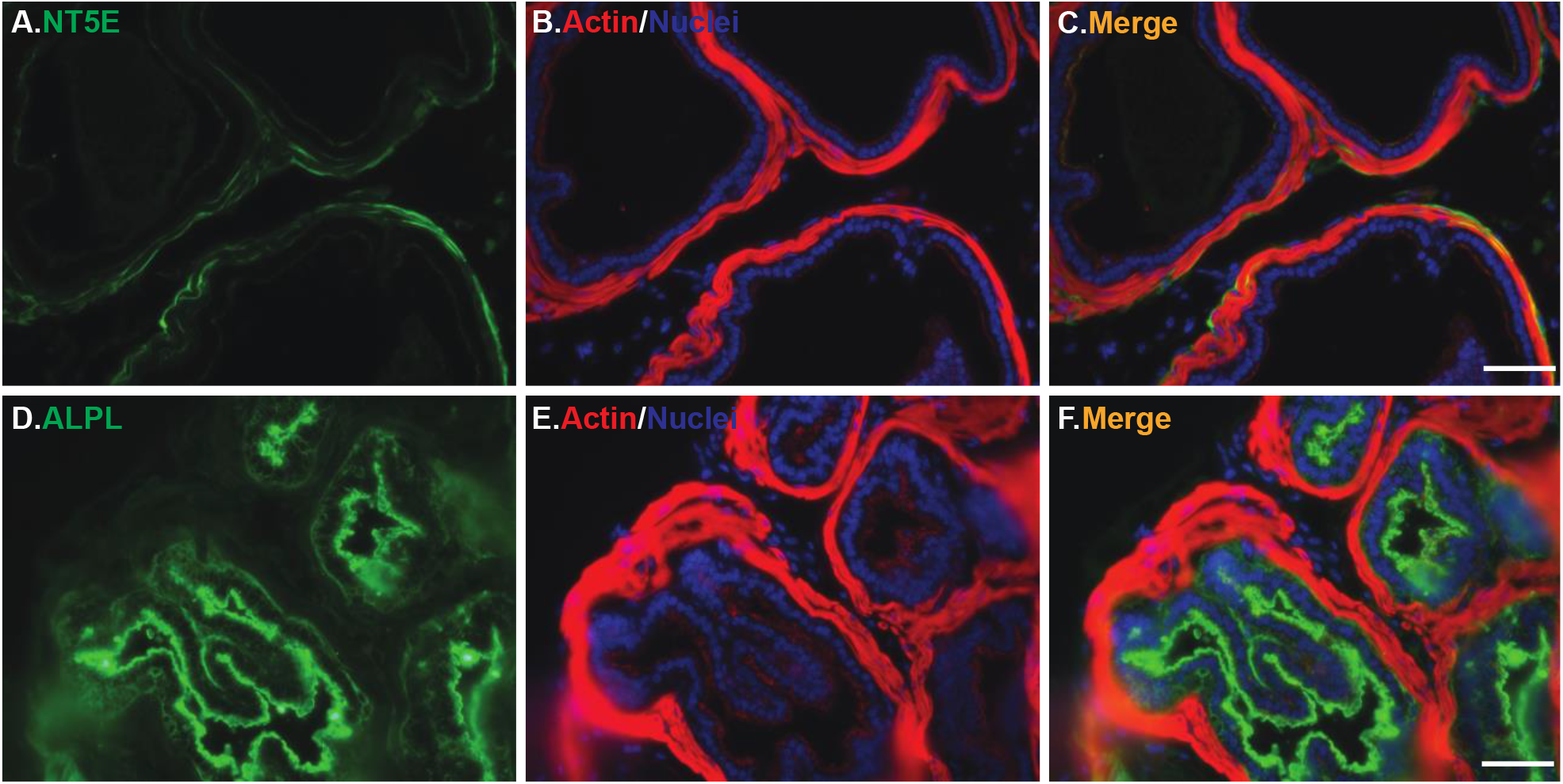
Expression of NT5E and ALPL in the prostate gland. Mouse prostate tissue was labeled with specific anti-NT5E (green in panel A) and anti-ALPL (green in panel D) antibodies. NT5E signaling did not colocalize with prostate smooth muscle (red in panel B) but mimicked the ENTPD2 expression pattern, indicating localization in interstitial cells. ALPL signaling was strongly present in the prostate epithelial cell membrane, particularly the apical membrane facing the lumen. Merged images are shown in panels C and F. Nuclei were stained with DAPI (blue). The white scale bar in the bottom right corner of panels C and F represents 50 μm.

### The mouse prostate gland exhibits purinergic contractility

We performed myography studies to determine whether the mouse prostate can respond to purinergic stimulation. As shown in Figure 5A&E, the mouse prostate generates a significant contraction force in response to 100 mM KCl-induced depolarization, with peak force reaching approximately 10mN. The mouse prostate also produces significant contraction force in response to electrical field stimulation (Figure 5B&E). The addition of 10μM α, β-meATP induced a noticeable contraction that decayed quickly (Figure 5C&E). This rapid decay is due to the activation and desensitization of P2×1 receptors in smooth muscle cells, a well-known characteristic of the P2×1 receptor[23]. However, after 10-15 minutes of α, β-meATP-induced desensitization, the subsequent addition of 25 μM ATPγS generated another noticeable contraction force (Figure 5D&E), suggesting that different receptor(s) mediate this purinergic contractility.

**Figure 5.**
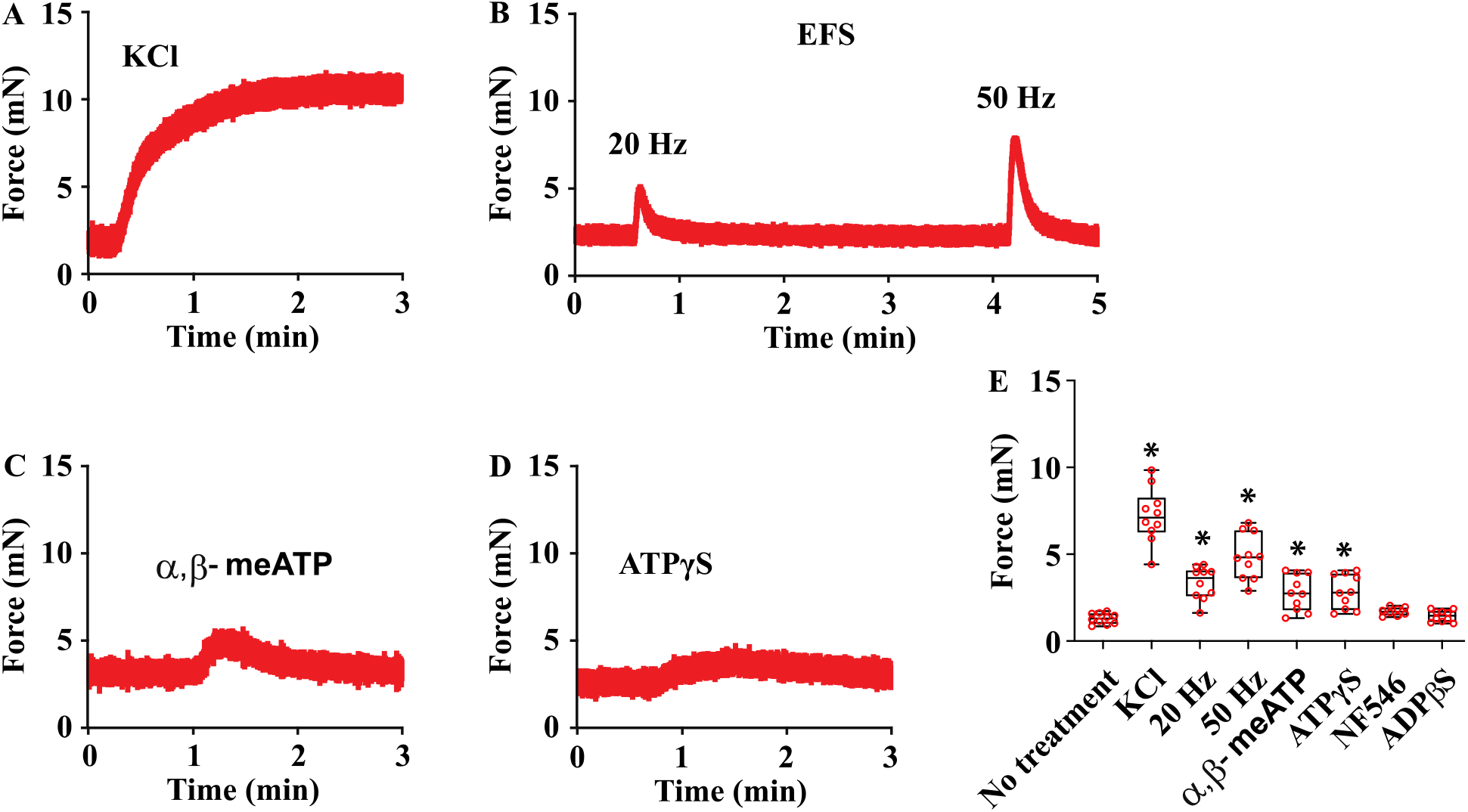
Purinergic contraction forces in mouse prostate tissue. (A-D) Representative traces of prostate smooth muscle contraction from male mice (n= 8-10 tissue preparations) in response to various stimuli: KCl depolarization, electrical field stimulation, P2X receptor agonist α, β-meATP, and P2 receptor agonist ATPγS. (E) Summary data presented as boxes-and-whiskers. The centerline represents the median, the box encompasses the interquartile range (IQR, 25th to 75th percentile), and the whiskers extend from the minimum to the maximum values.. Data were analyzed using the Student’s *t-*test. * p < 0.05.

In addition to activating P2X receptors, ATP also activates P2Y1, P2Y2, and P2Y11 receptors [24]. We tested NF546, a selective agonist for the P2Y11 receptor, but 15 μM NF546 produced no measurable response. We also tested the involvement of ADP-activated P2Y receptors by stimulating the prostate tissue with ADPβS, which yielded no observable contraction force (Figure 5E).

### P2Y receptors are potential candidate targets for novel purinergic contractile force

The observation of a novel ATPγS-induced contraction force suggests the potential presence of P2Y1, P2Y2, and P2Y11 receptors in the prostate gland. Consequently, we performed further immunofluorescent localization studies. Interestingly, all three ATP-activated P2Y receptors were detected in the prostate gland, specifically in the prostate smooth muscle cells (Figure 6A-I). This supports their potential role in mediating prostate smooth muscle contractility.

**Figure 6.**
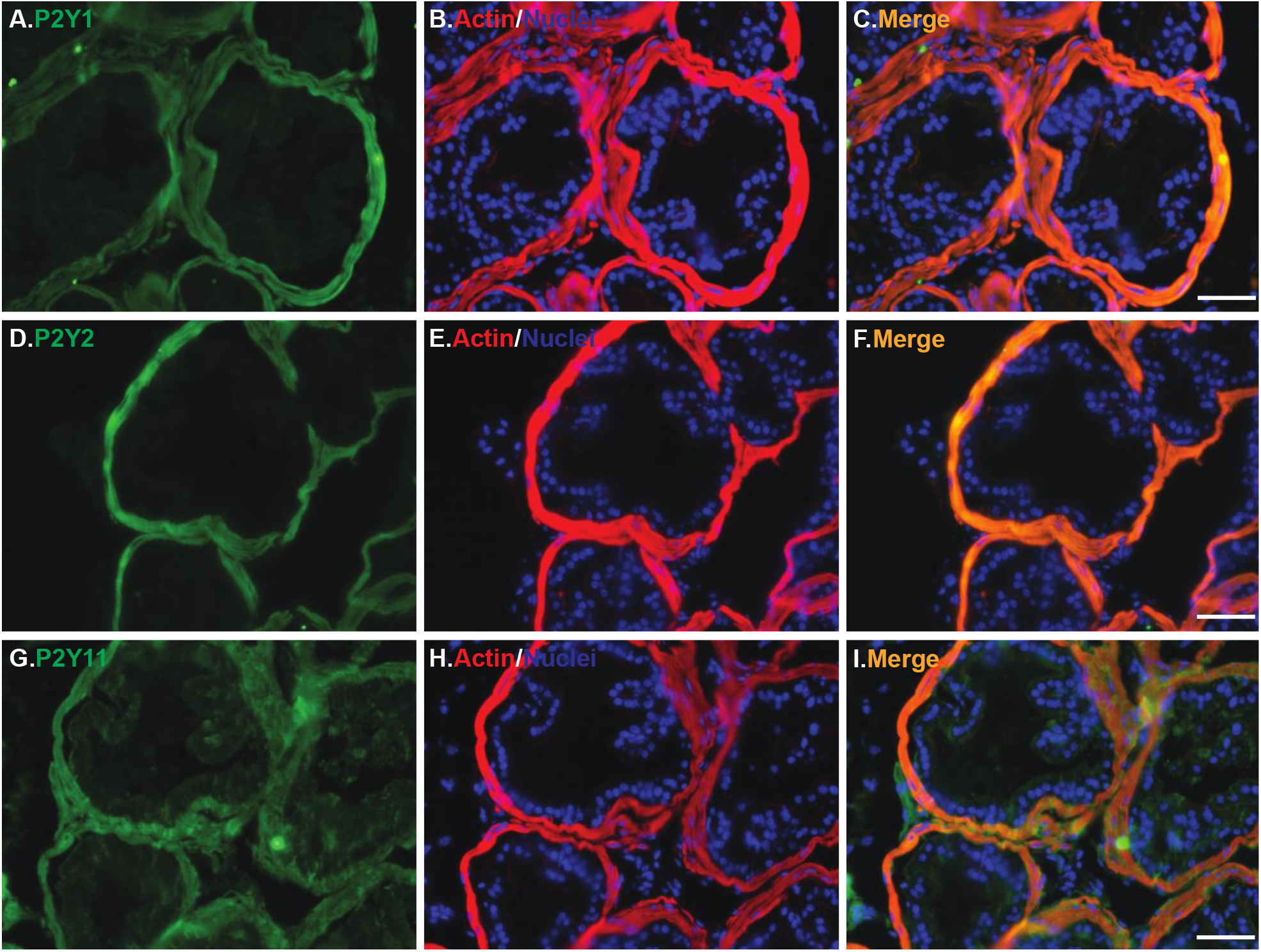
Expression of ATP-sensitive P2Y receptors in mouse prostate smooth muscle. (A, D, G): Mice prostate tissue labeled with specific antibodies against P2Y1 (green in panel A), P2Y2 (green in panel D), and P2Y11 (green in panel G) receptors. (B, E, H): Prostate smooth muscle labeled with a red marker. (C, F, I): Merged images showing colocalization of P2Y receptors (green) with prostate smooth muscle (red), resulting in yellow signals. Nuclei are stained blue with DAPI. The white scale bar in the bottom right corner of panels C, F and I represents 50 μm.

## Discussion

ATP is a well-characterized neurotransmitter but is also released from tissues due to mechanical stress, injury, or noxious stimuli. Excessive extracellular ATP can cause abnormal pain sensation, elevated muscle tone, and even tissue damage. Therefore, extracellular purine kinetics must be tightly regulated to maintain proper cellular functions[4]. For example, neuronally released ATP needs to be quickly recycled or degraded to avoid continuous activation of corresponding P2 receptors. This is achieved by cell surface purine-converting enzymes. Decreased purine hydrolysis activity has been reported in the prostates of aged mouse and guinea pig [9; 10].

Our data demonstrate the expression of multiple purine-converting enzymes in all three major types of prostate cells (Figures 1, 2, and 4). Although prostate smooth muscle cells exhibit strong ENTPD1 expression, they lack an enzyme to convert AMP to adenosine. However, prostate interstitial cells express both ENTPD2 and NT5e, allowing for the conversion of ATP/ADP to AMP and adenosine. Prostate epithelial cells show strong expression of ALPL, with intriguing strong apical expression, suggesting a role in controlling lumen ATP concentration in the prostate gland. The functionality of these differential expression patterns of purine-converting enzymes remains unclear but suggests an important role in prostate function.

There are very few reports on mouse prostate smooth muscle contractile function,likely due to its small size[9; 25]. Unlike the gastrointestinal or urinary bladder smooth muscle, which aligns longitudinally or circularly to allows optimized stretching of isolated muscle strips, prostate smooth muscle aligns irregularly around the glands. This anisotropy makes it impossible to stretch prostate smooth muscle on a tissue holder. Indeed, when prostate tissue was stretched, the tension increased quickly and did not decay significantly over time. In contrast, to the tension in gastrointestinal or urinary bladder smooth muscle tissue increases slowly and decays over time when stretched.

It is well known that smooth muscle stretch or length is a critical factor in generating maximal contraction force[26], while muscle with un-optimized length generates less force. Based on this theory, the contractile force of prostate muscle might be significantly smaller than that of gastrointestinal or urinary bladder smooth muscle tissue. This seems to be true, as the force generated by mouse prostate tissue in our study was significantly less than the reported forces generated by smooth muscle tissue in other systems [20; 27]. Nonetheless, we detected significant mouse prostate contraction in our experimental setting, with maximal force reaching approximately 10 mN in response to 100 mM KCl stimulation (Figure 5A).

The significance of purinergic contraction force in prostate smooth muscle remains debated in both rodents and humans. However, it is generally accepted that the P2×1 receptor is present in prostate smooth muscle and mediates ATP-induced contraction[6; 7; 8; 9; 10]. We have confirmed its presence in mouse prostate smooth muscle and demonstrate a noticeable contractile function in response to agonist 10μM α, β-meATP (Figures 1D-F & 5C). Interestingly, after P2×1 desensitization, a small but clear contraction force occurs in response to ATPγS, but not ADPβS (Figure D & E). This led us to identify multiple P2Y receptors in mouse prostate smooth muscle (Figure 6).

P2Y1 and P2Y2 receptors are coupled with Gq protein, which can activate phospholipase C and inositol triphosphate pathways, leading to smooth muscle contraction in the esophagus[28; 29]. Whether P2Y1 and P2Y2 receptors in mouse prostate smooth muscle have a similar function requires further investigation.

Smooth muscle plays an important role in prostate function by expelling seminal fluid. It is also an important drug target in BPH treatment by blocking the adrenergic alph1 receptor [30], which relaxes prostate smooth muscle tone. The increase in extracellular ATP and the decline in purine-converting enzyme activity in the aging prostate suggest a potential novel therapeutic approach for BPH. Further studies are needed to determine whether these purine-converting enzymes are also present in the human prostate and whether these novel P2Y receptors exist and are functional in both mouse and human prostates.

Limitations. The antibodies used in this study are mostly rigorously validated from previous reports, such as those on tissues from knockout animals. Although the antibodies for P2Y receptors are not rigorously validated, our studies produced clear and convincing results consistent with previous reports of these receptors in other systems[28; 29]. In addition to the purine-converting enzymes and P2 receptors studied, P1/adenosine receptors also play important roles in smooth muscle function, particularly in smooth muscle relaxation. However, based on our knowledge and experience, current anti-adenosine receptor antibodies are not reliable, and therefore, we did not attempt to localize these receptors.

In summary, we have identified multiple purine-converting enzymes in mouse prostate glands with differential localizations. Additionally, we have identified P2Y receptors in mouse smooth muscle that potentially mediate purinergic contractility. The presence of these molecules suggests that purinergic signaling may play a role in various prostate functions and dysfunctions, including inflammation, proliferation, metabolism, and muscle contraction.

## Author Contributions

J.Y. and C.S. performed tissue isolation, fixation, section, immunostaining, microscopy imaging, and myograph studies. Z.W. and A.O. contributed to the conception and interpretation of data. Z.W., A.O., and C.S. conceived and supervised the project. Z.W. and A.O. wrote the manuscript, and all authors critically discussed the results and reviewed the manuscript.

### Acknowledgments

The authors acknowledge fundings received from the National Institutes of Health/ National Institute on Diabetes and Digestive and Kidney Diseases grants 1R01DK124502,1R01DK140473,1R01DK142211.

## Competing interests

The authors declare no competing interests.

## Data availability Statement

The datasets used and analyzed during the current study are available from the corresponding author on reasonable request.

## Notes

### Competing Interest Statement

The authors have declared no competing interest.

